# Darwin’s bark spider shares a spidroin repertoire with *Caerostris extrusa* but achieves extraordinary silk toughness through gene expression

**DOI:** 10.1101/2021.07.16.452619

**Authors:** Nobuaki Kono, Rintaro Ohtoshi, Ali D Malay, Masaru Mori, Hiroyasu Masunaga, Yuki Yoshida, Hiroyuki Nakamura, Keiji Numata, Kazuharu Arakawa

## Abstract

Spider silk is a protein-based material whose toughness suggests possible novel applications. A particularly fascinating example of silk toughness is provided by Darwin’s bark spider (*Caerostris darwini*) found in Madagascar. This spider produces extraordinarily tough silk, with an average toughness of 350 MJ/m and over 50% extensibility, and can build river-bridging webs with a size of 2.8 m^2^. Recent studies have suggested that specific spidroins expressed in *C. darwini* are responsible for the mechanical properties of its silk. Therefore, a more comprehensive investigation of spidroin sequences, silk thread protein contents, and phylogenetic conservation among closely related species is required. Here, we conducted genomic, transcriptomic, and proteomic analyses of *C. darwini* and its close relative *Caerostris extrusa*. A variety of spidroins and low-molecular-weight proteins were found in the dragline silk of these species; all of the genes encoding these proteins were conserved in both genomes, but their genes were more expressed in *C. darwini*. The potential to produce very tough silk is common in the genus *Caerostris*, and our results may suggest the existence of plasticity allowing silk mechanical properties to be changed by optimizing related gene expression in response to the environment.

## Introduction

Since its discovery in Madagascar in 2010, Darwin’s bark spider, *Caerostris darwini* (Araneae: Araneidae), has fascinated the world because of its unique behaviour, ecology, and biomaterial production ability and has become a model organism for addressing the evolution of environmental adaptation in spiders ^1^. Darwin’s bark spider is an orb-weaving spider capable of creating large river-bridging webs with anchor threads as long as 25 m and sizes of up to 2.8 m^2 1,2^. The size of the webs built by two other species, *Nephilengys borbonica* and *Trichonephila inaurata*, producing relatively large orb webs reaches only approximately 0.4 m^2^, indicating the exceptional size of the river-bridging webs of *C. darwini* ^3,4^. It is thought that its extremely large orb web remains stable and unbroken because it consists of tough dragline silk spun from a relatively long spinning duct of the major ampullate gland ^5^. Spider silk is a typical high-performance protein material and with the attractive properties of high toughness, strength and extensibility ^6^. The silk of Darwin’s bark spider, in particular, shows an extraordinarily high level of toughness. A previous study on the mechanical properties of Darwin’s bark spider silk reported tensile strength, extensibility, and toughness values of 1.6 GPa, 52%, and 354 MJ/m^3^, respectively ^4^. Other orb-weaving spiders, such as *Trichonephila clavipes* or *Nephila pilipes*, show silk toughness ranging from only 100 – 150 MJ/m^3^ on average ^7^. Some novel specific amino acid motifs in spidroins have been suggested as candidate factors responsible for the excellent mechanical properties of *C. darwini* silk.

A previous transcriptome analysis of Darwin’s bark spider revealed that major ampullate spidroin (MaSp) families 4 (MaSp4) and 5 (MaSp5) harbour unique motifs ^5,8^. Among these homologous groups, MaSp is a main component of dragline silk and is encoded by at least five gene families. MaSp families 1 and 2 (MaSp1 and MaSp2) have been well studied over a long period, whereas MaSp family 3 (MaSp3) was revealed by recent genomic and transcriptomic analyses ^9,10,11^. MaSp4 and MaSp5 are new homologs that have so far only been found in Darwin’s bark spider ^5^. MaSp4 and MaSp5 contain “VSVVSTTVS” and “GGLGGSG” motifs, respectively, specific to Darwin’s bark spider in their repetitive domains, and these proteins have been reported as possible factors contributing to silk toughness ^5^. However, the sequence architectures of MaSp4 and MaSp5 have been only partially identified, and their detailed variation and conservation and even their actual role in dragline silks remain unclear. Knowledge of gene sizes, motif patterns, and phylogenetic conservation is essential for discussing the evolution of silk mechanical properties. In addition, it has been recently elucidated that spider silk is made of composite materials and contains various proteins other than spidroins. Spider silk-constituting elements (SpiCEs) are non-spidroin low-molecular-weight (LMW) proteins of unknown function that have been widely found in the family Araneidae and were recently investigated to determine their effects on the formation and mechanical properties of spider silk ^9,12,13,14,15^.

Here, to enable comparative analysis between closely related species, we present two bark spider draft genomes and conduct a multiomics analysis. As a close relative of Darwin’s bark spider, we use *Caerostris extrusa. C. extrusa* is clearly defined as a different species even though it lives in the same region of Madagascar because the genetic distances between individuals of these species are much larger than those within the species inferred from DNA barcodes ^16^. Using draft genomes prepared by applying hybrid sequencing technology, we conducted a multiomics analysis to curate a highly accurate spidroin catalogue, conduct phylogenetic searches of spidroins, and profile their protein and mRNA expression. Based on these analyses, we describe candidate elements contributing to the differences in silk mechanical properties according to the identified genes or their expression patterns.

## Results

### Silk comparison between bark spiders

To understand the uniqueness of Darwin’s bark spider (*C. darwini*) silk is among known spider silks, we reeled dragline silks from two bark spiders (*C. darwini* and *C. extrusa*) and measured the mechanical properties and wide-angle X-ray scattering (WAXS) profiles of the reeled dragline silks. The diameter of *C. darwini* dragline silk was on average 2.3 times larger than that of *C. extrusa*, and the crystallinity of *C. darwini* silk was 30%, compared to the 23% crystallinity of *C. extrusa* silk (Fig. 1a-d). Tensile tests confirmed the extraordinary toughness of Darwin’s bark spider silk, as reported in a previous study ^4^. The toughness of *C. darwini* silk was approximately twice that of *C. extrusa* silk, and the parameter that contributed most to this difference was extensibility (Fig. 1e, f). The extensibility of *C. extrusa* dragline silk was approximately 23.5%, which is an average value within the family Araneidae ^17^, while that of *C. darwini* was up to 49.73% (Table 1). It is particularly interesting that this dragline silk can maintain such high extensibility while maintaining a tensile strength above 1 GPa. A comparative multiomics analysis was then carried out to investigate how *C. darwini* could produce such extraordinarily tough silk.

**Table 1:**
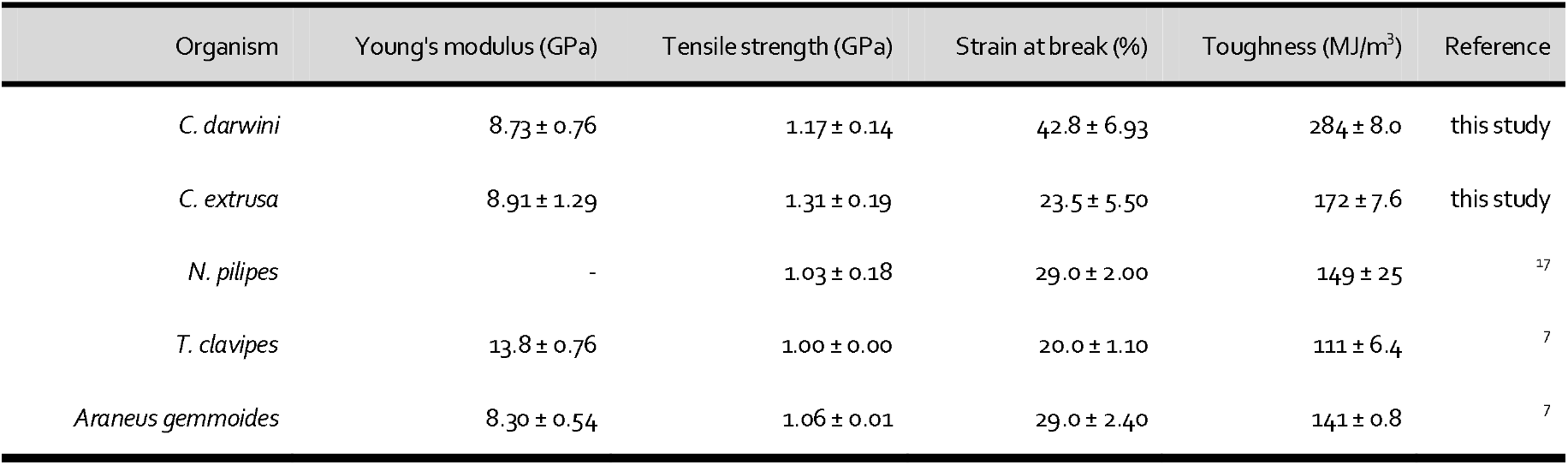
Mechanical properties of orb-weaving spider silks

**Fig. 1:**
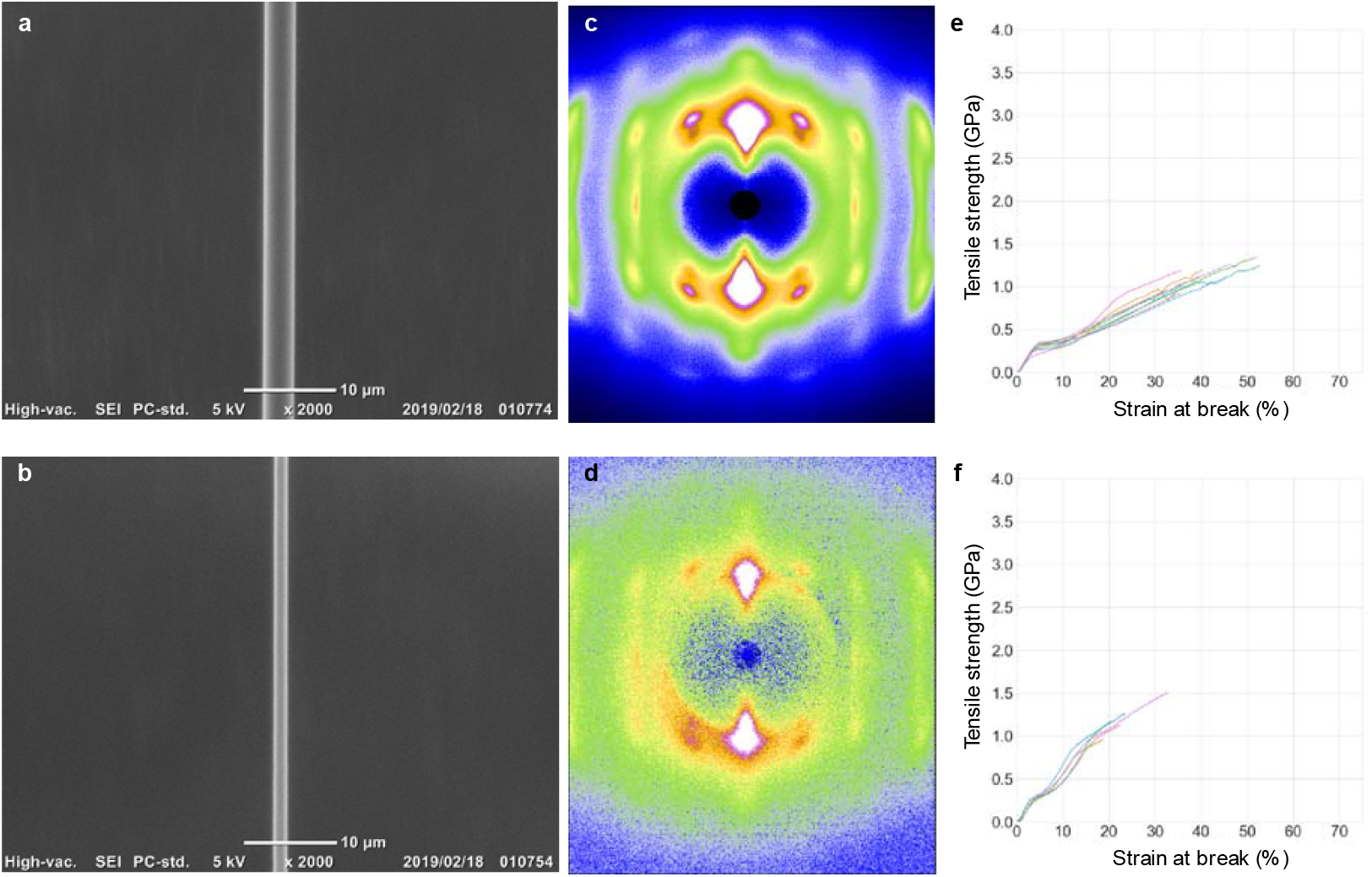
Structures and mechanical properties of dragline silks. **a, b** Scanning electron micrographs (SEMs), **c, d** 2D WAXS pattern images, and **e, f** stress-strain (S-S) curves of *C. darwini (***a, c, e**) and *C. extrusa* (**b, d, f**). Scale bar in **a** and **b** is 10 µm. S-S curves are the results of tensile tests of dragline silks from *C. darwini* (toughness: 284 ± 80 MJ/m^3^ ; tensile strength 1.17 ± 0.14 GPa; strain at break: 42.8 ± 6.93%) and *C. extrusa* (toughness: 172 ± 76 MJ/m^3^ ; tensile strength: 1.31 ± 0.19 GPa; strain at break: 23.5 ± 5.5%).

### Genome sequences of bark spiders

We present the draft genome sequences of two bark spiders (*C. darwini* and *C. extrusa*) (Fig. 2a, b). The de novo sequencing of these large, complex spider genomes is challenging, and we sequenced the bark spider genomes via hybrid sequencing with a combination of Nanopore, 10x GemCode, and Illumina technologies. Genomic DNA (gDNA) was extracted from dissected legs of adult female individuals. The Nanopore gDNA sequencing of the produced 10.23 million long reads with an N50 length of over 5.34 kbp. In *C. darwini*, in addition to long reads, 990 million GemCode-barcoded 150-bp paired-end reads were sequenced by using Illumina technology. These gDNA sequenced reads were assembled, and the numbers of scaffolds (and N50 lengths) in the *C. darwini* and *C. extrusa* draft genomes were 15,733 (and 440,877 bp) and 21,729 (and 98,474 bp), respectively (Fig. 2c, d, and Table 2). The genome size of *C. darwini* was estimated with GenomeScop ^18^ based on the k-mer distribution to be 1.58 Gbp, which is relatively small relative to other spider genomes, which average over 2.50 Gbp in size ^19^. The assembled scaffolds were assessed by Benchmarking Universal Single-Copy Ortholog (BUSCO) analysis ^20^, and the completeness scores of *C. darwini* and *C. extrusa* were 92.6% and 82.7%, respectively (Fig. 2c, d, and Table 2).

**Table 2:**
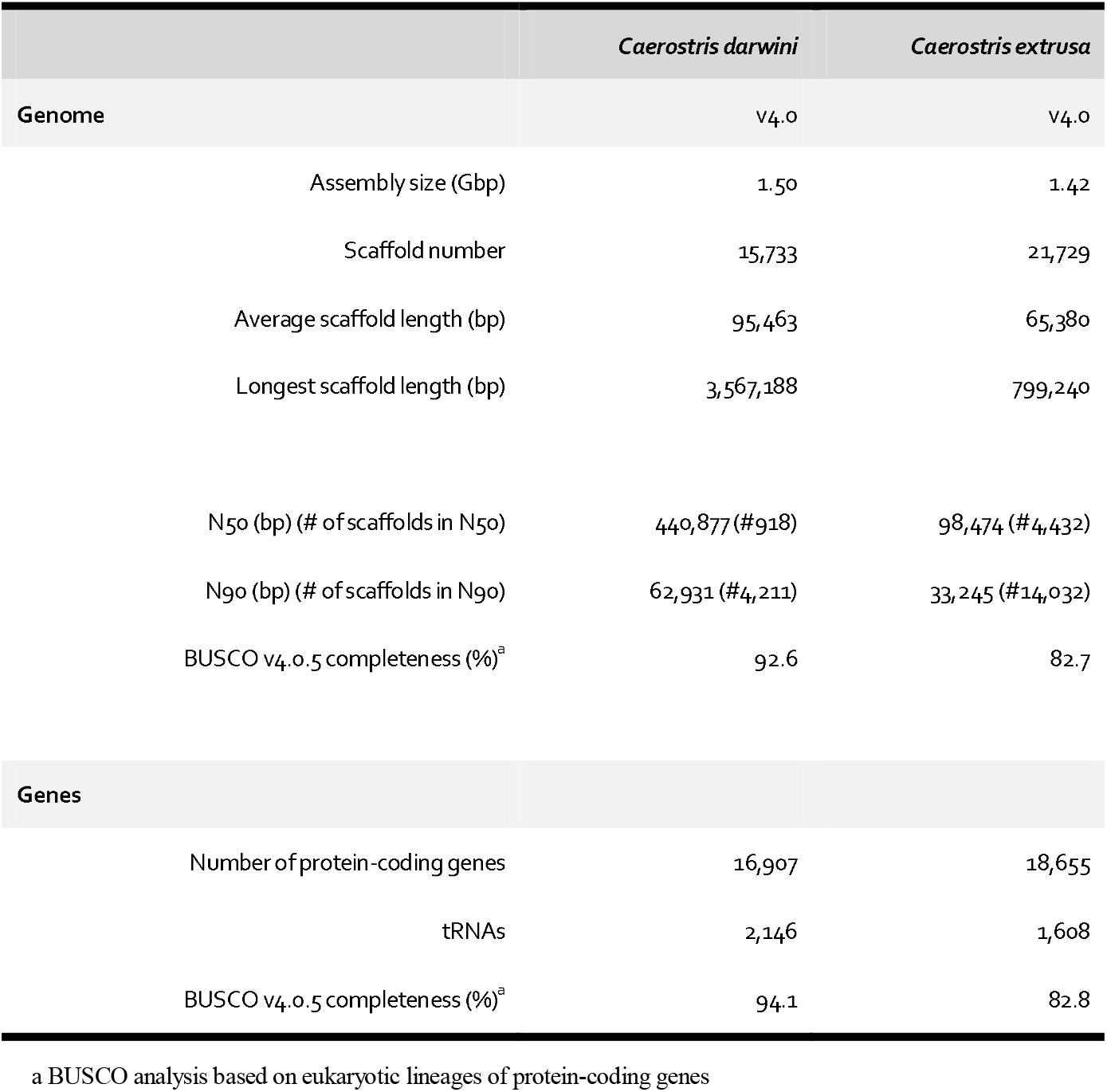
Genome statistics

|  | *Caerostris darwini* | *Caerostris extrusa* |
| --- | --- | --- |
| **Genome** | v4.0 | v4.0 |
| Assembly size (Gbp) | 1.50 | 1.42 |
| Scaffold number | 15,733 | 21,729 |
| Average scaffold length (bp) | 95,463 | 65,380 |
| Longest scaffold length (bp) | 3,567,188 | 799,240 |
| N50 (bp) (# of scaffolds in N50) | 440,877 (#918) | 98,474 (#4,432) |
| N90 (bp) (# of scaffolds in N90) | 62,931 (#4,211) | 33,245 (#14,032) |
| BUSCO v4.0.5 completeness (%)<sup>a</sup> | 92.6 | 82.7 |
| **Genes** |  |  |
| Number of protein-coding genes | 16,907 | 18,655 |
| tRNAs | 2,146 | 1,608 |
| BUSCO v4.0.5 completeness (%)<sup>a</sup> | 94.1 | 82.8 |

<sup>a</sup> BUSCO analysis based on eukaryotic lineages of protein-coding genes

**Fig. 2.**
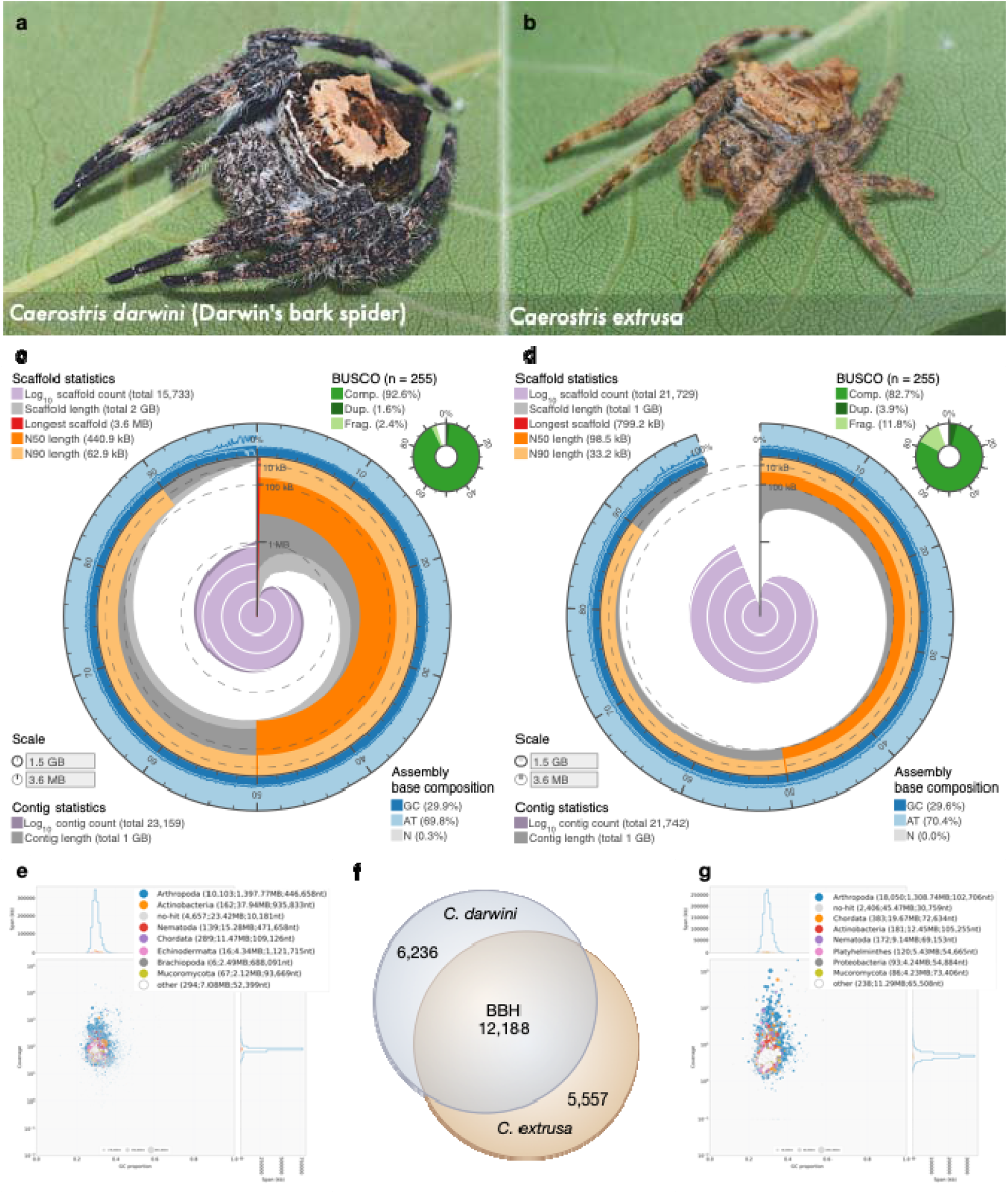
Overview of two bark spiders. Spider images and genome statistics of *C. darwini* (**a, c, e**) and *C. extrusa* (**b, d, g**). **c** and **d** are the assembly and genome statistics with BUSCO completeness. **e** and **g** are the results from the contamination check using BlobTools. These plots show the taxonomic affiliation at the phylum rank level, distributed according to GC% and coverage. **f** The number of orthologous genes identified by bidirectional best hits (BBH, 1.0e-5) between expressed gene groups (TPM > 1.0).

Gene prediction was conducted using a gene model generated from cDNA-seq mapping data. The cDNA library was constructed using mRNA extracted from abdomen samples of each species, and Illumina sequencing produced approximately 35 million reads per sample. The numbers of protein-coding genes initially predicted were 56,046 and 82,821 for *C. darwini* and *C. extrusa*, respectively. Redundant genes were eliminated based on identity, expression levels, and annotation, and 16,907 and 18,655 functional protein-coding gene sets with BUSCO completeness scores of 94.1% and 82.8% were obtained from *C. darwini* and *C. extrusa*, respectively (Table 2). The number of orthologous genes identified by bidirectional best hits (BBH, 1.0e-5) between expressed gene groups (TPM > 1.0) was 12,188 (Fig. 2f).

### Full spidroin catalogues

In addition to the known classical spidroins, various families or subfamilies specific to the genus *Caerostris* were all found to be shared between the two newly obtained genomes (Fig. 3). Each spidroin was assigned a name according to the previously reported nomenclature ^5,10,11,15^. Five families of MaSp, the main component of dragline silk, were observed, and nine genes (including paralogues) encoding these proteins were found. The full-length sequences were determined for almost all MaSp genes so that each gene type could be strictly distinguished. The repetitive units and gene lengths varied from 4-10 kbp, while the terminal domains were well conserved.

**Fig. 3.**
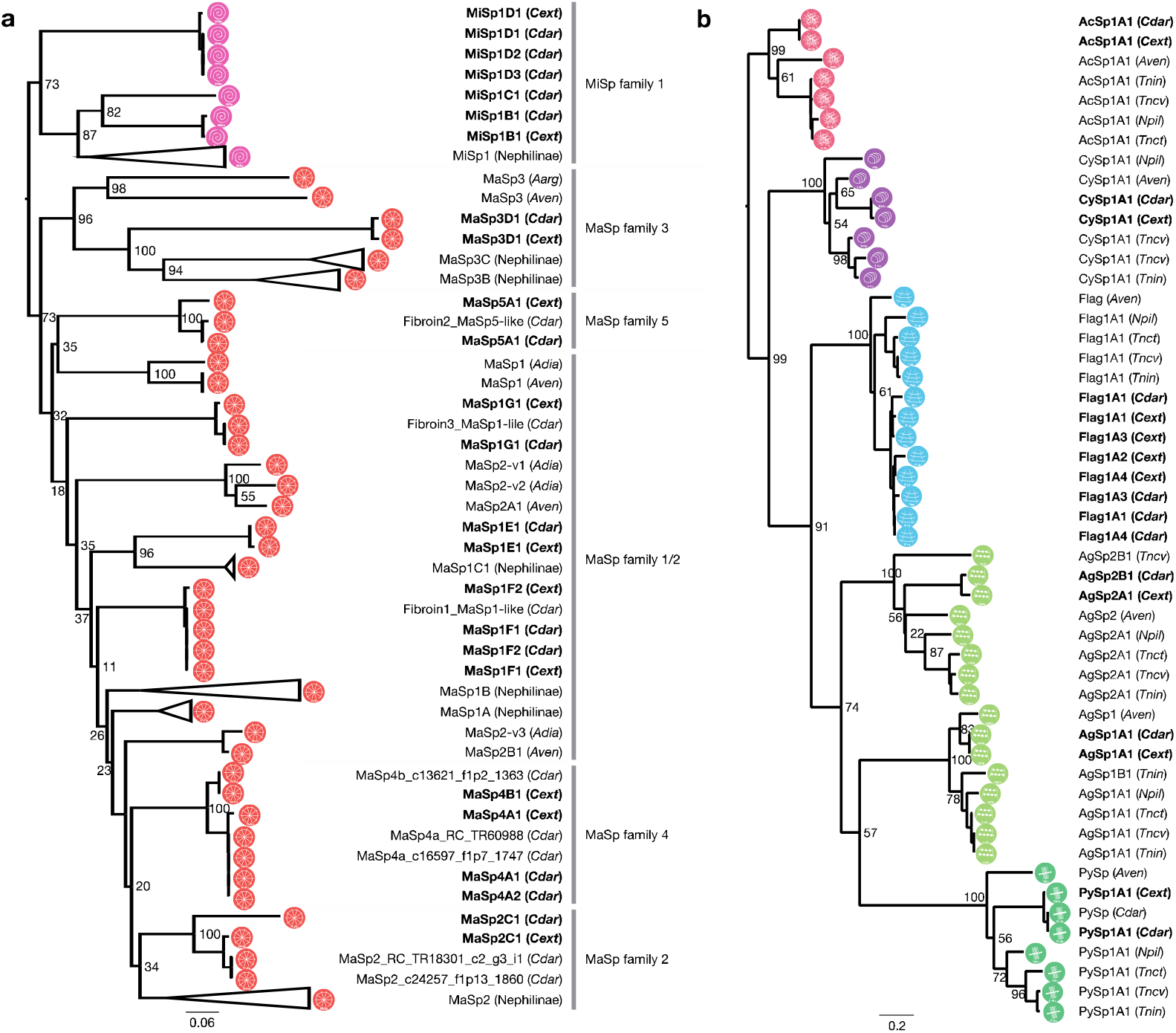
Phylogenetic tree of orb-weaving spider spidroins. Phylogenetic analysis of all spidroins (**a** ampullate spidroins and **b** other spidroins) with 100 aa N-terminal regions (*Cdar*: *C. darwini, Cext*: *C. extrusa, Aarg*: *Argiope argentata, Aven*: *A. ventricosus, Adia*: *Araneus diadematus, Tnct*: *T. clavata, Tncv*: *T. clavipes, Tnin*: *T. inaurata madagascariensis, Npil*: *N. pilipes*). Genus *Caerostris* spidroins are indicated by bold font. Branch labels are bootstrap support values.

MaSp4 and MaSp5, reported in a previous study ^5^, was found in both bark spiders. The GPGPQ repetitive motifs of MaSp4 were unique and differed from those of the spidroins reported thus far. However, a phylogenetic analysis showed that all of the MaSp4s belonged to the same clade as the MaSp2 proteins based on their terminal domains, suggesting that they may constitute a lineage of MaSp2 (Fig. 3a and Supplementary Fig. 1a, b). This result supported the previous study ^5^.

In general, GPGQQ is considered the MaSp2-specific motif in the family Araneidae ^21^, but the bark spider MaSp2s had GPGSQ motifs in which the diglutamine (QQ) was replaced by SQ (Table 3 and Supplementary Fig. 1c). Therefore, the GPGPQ motif of MaSp4 can be considered an alternative to GPGQQ. In addition, MaSp4 did not contain a poly-A sequence, like MaSp2, but instead contained a VSVVSTTVS sequence, composed of neutral amino acids other than alanine (Table 3 and Supplementary Fig. 1c). Hence, if SQ or PQ and poly-X (X=neutral amino acids) sequences are viable alternatives to QQ and poly-A sequences, it may be reasonable to consider MaSp4 as a new subfamily of MaSp2.

**Table 3:**
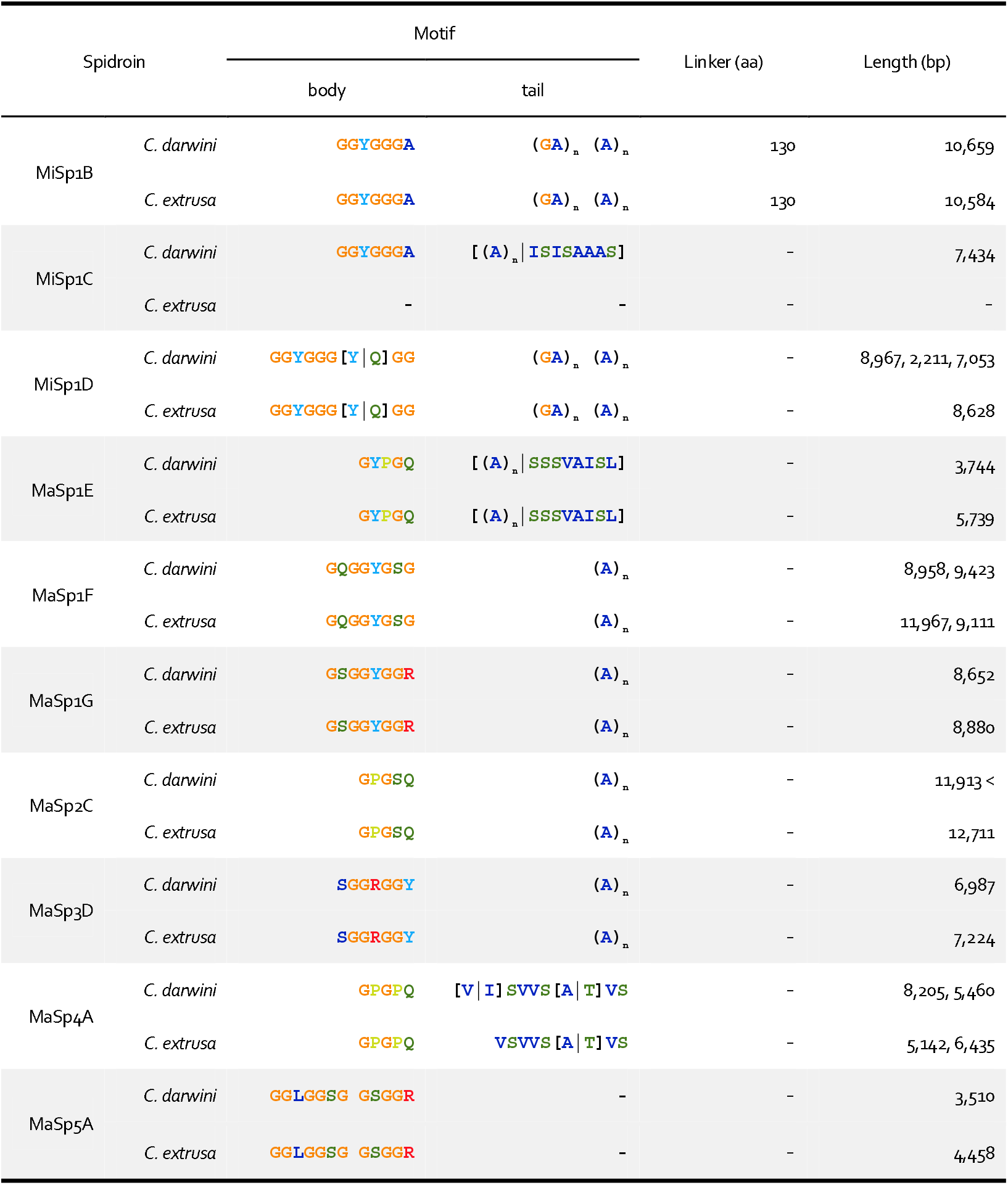
Repeat motifs in *Caerostris* MiSp and MaSp

The full-length of MaSp5 was also revealed. The size of MaSp5 was 3.5 kbp, which is relatively small among spidroins. The MaSp5 proteins did not contain a tail portion of the repetitive unit (such as the poly-A sequence present in other MaSp families) and only showed tandemly arranged GGLGGSG or GSGGR motifs (Table 3). The MaSp5 N-terminal regions domain did not cluster into the same clade as those of any other MaSp family; thus, MaSp5 seems to be a new family of MaSp, as suggested by previous studies ^5^. In addition, we confirmed the presence of MaSp family 3 (MaSp3) in the two bark spider genomes by phylogenetic analysis (Fig. 3a). The MaSp3 proteins of the Araneidae family contain a DGGRGGY motif ^9,10^, but the MaSp3s found in the bark spiders contained a SGGRGGY motif, in which aspartic acid (D) was replaced by serine (S) (Table 3).

No significant differences in spidroin other than MaSp were observed between the two species. A total of five minor ampullate spidroins (MiSps) were identified and classified into three subfamilies (MiSp1B, MiSp1C, and MiSp1D). MiSp1B and MiSp1D had typical MiSp motif tails (poly-A/GA), but MiSp1C had an alternative poly-X tail, as observed in MaSp4. Multiple spidroin paralogues were also observed for flagelliform spidroin (Flag), the main component of the core fibre of the prey capture thread, including five paralogues in the *C. darwini* genome and six in the *C. extrusa* genome (Fig. 3b and Supplementary Fig. 2).

These spidroin characteristics and types were conserved in the two bark spiders without exception. Therefore, since omics approaches beyond the genome level were required to explain the toughness of *C. darwini* dragline silk, we performed protein and mRNA expression profiling.

### Expression profiling of proteins in dragline silk and mRNAs in the spider body

Proteome analysis was performed with nanoElute and timsTOF using dragline silks reeled from adult female spiders. The obtained spectra were annotated based on our draft genome database, and dozens of proteins were detected. The dragline silk contained all MaSp families 1-5, with MaSps alone accounting for more than 80% of the total proteins (Fig. 4a). The proteome analysis of the dragline silk also demonstrated the presence of a small percentage of non-spidroins of unknown function. We defined the top four non-spidroin genes whose expression was confirmed by transcriptome analysis of the genus *Caerostris* dragline silk as SpiCEs, which were designated SpiCE-CMa1 to 4. SpiCEs are defined as non-spidroin LMW proteins associated with spider silk, of unknown function and showing both mRNA and protein expression ^9^. Among these proteins, those with a high cysteine content are known as cysteine-rich proteins (CRPs) ^14^. SpiCE-CMa4 contained 10% cysteine, suggesting that it was a CRP member (Fig. 4a). It was also notable sequence feature that SpiCE-CMa3 is rich in glycine and other hydrophobic amino acids. These SpiCE gene sequences were well conserved in the genomes of the two bark spiders, as were their spidroin sequences (Fig. 4b). However, a partial deletion of the disordered region was observed only in the gene sequence of *C. extrusa* SpiCE-CMa3. On average, the four SpiCEs contained 1% to 5% of dragline silk and SpiCE-CMa3 was the most abundant protein. Furthermore, the expression levels of several genes in dragline silk were observed to differ between the two spiders. According to the results of mRNA expression profiling by transcriptome analysis, the *MaSp4* and *MaSp5* genes, which were found to be *Caerostris* specific, were expressed at levels 2-6 times higher in *C. darwini* than in *C. extrusa*. In addition, the expression levels of almost all SpiCEs were more than 2-fold higher in *C. darwini* than in *C. extrusa* (Fig. 4c).

**Fig. 4:**
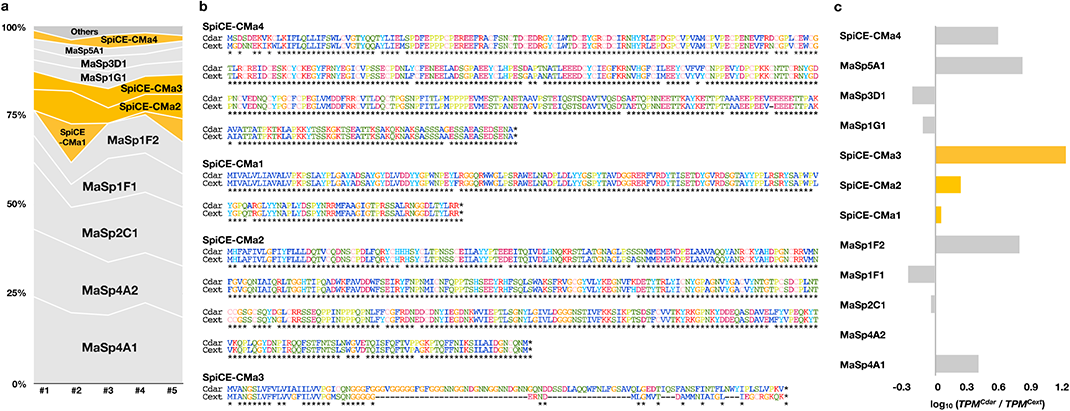
Expression profiles of protein and mRNA. **a** The results of the proteome analysis of five dragline silk samples. Spidroins and SpiCEs are indicated by grey and orange. **b** Sequence alignment of SpiCEs from *C. darwini* and *C. extrusa*. **c** Gene expression ratio between *C. darwini* and *C. extrusa*. Spidroins and SpiCEs are indicated by grey and orange, respectively.

## Discussion

Genome information allows comprehensive gene prediction and annotation and aids in large-scale transcriptomic, proteomic, and phylogenomic analyses. However, in spider research, the available molecular information is limited in many cases. One of the main reasons for this is the large genome sizes of spiders. We have solved this problem by using hybrid sequence technology and obtained two bark spider genomes. The genome sizes of the bark spiders were smaller than those of other Araneidae spiders (approximately 3 Gbp), but they were still more than 1 Gbp, in line with the size of the *Argiope bruennichi* genome ^22^. The Nanopore long reads contributed significantly to the construction of a highly accurate spidroin catalogue.

Our catalogue confirmed the existence of *Caerostris*-specific MaSp paralogues, namely, MaSp families 4 and 5, reported in a previous study ^5^. According to the full-length sequence of MaSp4 identified in this work, the gene previously reported as MaSp4 was clearly shown to cluster with the MaSp2 family based on the sequence similarity of terminal amino acid motifs (Fig. 3). The repetitive motifs also showed resemblance to the MaSp2 family. The previously identified QQ of *Caerostris* MaSp2 was found to be substitutable with SQ in the genus *Caerostris* (Table 3); PQ was likewise used substitutably in MaSp4. The polyalanine region was replaced with other neutral amino acids (S, V, I, or T). Similar replacement of polyalanine motifs with other neutral amino acids was observed in the MaSp family 1 (MaSp1), in which the tail portion of MaSp1E or MaSp1C repetitive unit contained a mixture of poly-A and SSSVAISL or ISISAAAS motifs, respectively (Table 3). Overall, extreme specialization of the amino acid motifs of MaSp paralogues seems to have occurred in the *Caerostris* genus. Previous studies of the genomes of *Araneus ventricosus* and *Trichonephila clavata* identified the existence of the Araneoidea-specific MaSp3 family of spidroins as well as the clade-specific non-spidroin component SpiCE proteins, both of which are essential for the high mechanical performance of silks in these species ^9,15^. *Caerostris*, classified in the family Araneidae, also possesses MaSp3 paralogues as well as clade-specific SpiCEs.

Intriguingly, all of these genes were shown to be conserved in both *C. darwini* and *C. extrusa*, and these two species showed little difference in terms of the silk gene repertoire, although the mechanical performance of the silks of these two species differed significantly.

What then, are the elements contributing the toughness of Darwin’s bark spider silk? Our expression analysis showed differences in the deployment of the gene repertoire. *C. extrusa* showed a typical expression profile of Araneoidea, with high expression levels of MaSp3 and canonical MaSp1 and MaSp2 paralogues ^5,9,11,15,23^. Conversely, the expression of the *Caerostris*-specific paralogues *MaSp4* and *MaSp5* genes was greatly increased by 2-6 times in *C. darwini*. Furthermore, the expression of SpiCE-CMa3 was almost doubled in *C. darwini*, and the length of SpiCE-CMa3 in *C. extrusa* was approximately half that in *C. darwini* (Fig. 4a). These changes together may provide one explanation for the significant differences in the mechanical performance of the draglines of these two species.

Previous proteomics analyses of *A. ventricosus* and *T. clavata* silks showed a predominance of MaSp3 in the silk composition of these species ^9,15^. However, the proteome analysis of *C. darwini* dragline silk demonstrated that the proteins of the MaSp2 class (MaSp2 and MaSp4) were its major components, accounting for approximately 50% of the total proteins content. As shown in Fig. 1, the high toughness of *C. darwini* silk is mainly accounted for by enhanced extensibility. The presence of spidroins with many prolines in the repetitive domain (as found MaSp2) is known to increase the extensibility of spider silk ^24,25,26,27,28,29^, and the predominance of the MaSp2 class in the dragline silk of *C. darwini* therefore seemed to be in accord with its high extensibility (Fig. 4), explaining the long stretch beyond the yield point in particular (Fig. 1e). The initial stretching of the spider silk reaches the yield point with the rupture of the hydrogen bonds in semiamorphous regions (helices and β-turns), and subsequent deformation involves the β-sheets in the crystal domain ^30^. The spidroin β-sheets are composted of poly-A motifs ^26,31,32,33,34^. Johansson and Rising have proposed a hypothesis about the implications of the replacement of poly-A sequences with other amino acids for silk engineering ^35^ ; for example, alanine polymers are proposed to be replaceable by polymers of valine (poly-V) or isoleucine (poly-I) to form stiff sheets ^36^. The bark spider sequences do not contain valine and isoleucine as homopolymers, but they are included in combination with other amino acids, as in the ISVVSTTVS motif (Table 3). This semiconservative mode of replacement, accompanied by expression level regulation, may improve β-stacking after the yield point to a greater extent than is found in other silks.

In summary, the differences in the mechanical properties of silks between the two studied bark spiders were attributed to differences in the predominant spidroins. Although the existence of the clade-specific MaSp paralogs MaSp4 and MaSp5 in *C. darwini*, previously identified by transcriptome analysis, was confirmed by our genome analysis, these genes are also conserved in *C. extrusa*, whose silk does not show comparable performance. However, differences in the deployment of the shared repertoire of silk genes in these two species may be the key to the differences in their silk, as the predominant use of MaSp2 family proteins, including MaSp4s, in *C. darwini* is in accord with the significant enhancement of elasticity in its dragline. Research on the detailed contributions of these components will help to further explain the molecular mechanism underlying the extraordinary toughness of *C. darwini* silk.

## Methods

### Sample collection

Adult female spiders (*C. darwini* and *C. extrusa*) were collected from Andasibe, Eastern Madagascar (18°56’49.3’’S, 48°25’09.1’’E) and Parc National Andasibe Mantadia, Eastern Madagascar (18°56’08.3’’S 48°24’53.2’’E). The spiders were identified based on morphological characteristics and cytochrome c oxidase subunit 1 (*COI*) sequences in the Barcode of Life Data System (BOLD: http://www.barcodinglife.org). The natural dragline silks used for all experiments were sampled directly from adult female bark spiders restrained with two sponge pieces and locked with rubber bands. Silk reeling was performed at a constant speed (1.28 m/m for 1 h) with a reeling device developed by Spiber Inc. The specimens collected according to a previously established field sampling method ^37^ were transported to the laboratory, immersed in liquid nitrogen (LN2), and stored at -80 °C until subsequent processing. gDNA and total RNA were extracted from the samples.

### High-molecular-weight (HMW) gDNA extraction and genome sequencing

#### Extraction, purification, quality-quantification

HMW gDNA was extracted from the legs of flash-frozen spiders using Genomic-tips 20/G (QIAGEN) based on previous studies ^9^. The specimens were gently and quickly homogenized using a BioMasher II homogenizer (Funakoshi) and mixed with 2 mL of Buffer G2 (QIAGEN), including 200 µg/mL RNase A and 50 µL proteinase K (20 mg/mL). After incubation at 50 °C for 12 h on a shaker (300 rpm), the mixed lysate was centrifuged at 5,000 x g for 5 min at 4 °C, and the aqueous phase was loaded onto a pre-equilibrated QIAGEN Genomic-tip 20/G by gravity flow and washed three times. The DNA was eluted with a high-salt buffer (Buffer QF) (QIAGEN), desalted and concentrated using isopropanol precipitation and resuspended in 10 mM Tris-HCl (pH 8.5). The extracted gDNA was qualified using a TapeStation 2200 instrument with genomic DNA Screen Tape (Agilent Technologies) and quantified using a Qubit Broad Range dsDNA assay (Life Technologies). The purified gDNA was size-selected (> 10 kb) with a BluePippin with High Pass Plus Gel Cassette (Sage Science).

#### Library preparation and sequencing

Nanopore library preparation was implemented following the 1D library protocol (SQK-LSK109, Oxford Nanopore Technologies). The quality of the prepared library was calculated by the TapeStation 2200 system with D1000 Screen Tape (Agilent Technologies). Sequencing was performed using a GridION instrument with a SpotOn Flow Cell Rev D (FLO-MIN106D, Oxford Nanopore Technologies). Base calling was performed after the runs with Guppy base calling software (version 3.2.10+aabd4ec). For 10X GemCode library preparation, purified gDNA fragments longer than 60 kb (10 ng) were used to prepare the library with a Chromium instrument and Genome Reagent Kit v2 (10X Genomics) following the manufacturer’s protocol. 10X GemCode library sequencing was conducted with a NextSeq 500 instrument (Illumina) using 150-bp paired-end reads with a NextSeq 500 High Output Kit (300 cycles).

### RNA extraction and cDNA sequencing

RNA extraction was implemented based on a spider transcriptome protocol ^37^. Flash-frozen dissected abdomen tissue was immersed in 1 mL TRIzol Reagent (Invitrogen) along with a metal cone and homogenized with a Multi-Beads Shocker (Yasui Kikai). After the addition of chloroform, the upper aqueous phase containing RNA was automatically purified with an RNeasy Plus Mini Kit (QIAGEN) on a QIAcube instrument (QIAGEN). The quantity and quality of the purified total RNA were calculated with a Qubit Broad Range RNA assay (Life Technologies) and a NanoDrop 2000 system (Thermo Fisher Scientific). The RNA integrity number (RIN) was estimated by electrophoresis using a TapeStation 2200 instrument with RNA ScreenTape (Agilent Technologies). mRNA was selected from the total RNA using oligo d(T). cDNA was synthesized from mRNA isolated from 100 µg of total RNA by NEBNext Oligo d(T)25 beads (skipping the Tris buffer wash step). First- and second-strand cDNA were synthesized using ProtoScript II Reverse Transcriptase and NEBNext Second Strand Synthesis Enzyme Mix. cDNA library preparation was performed according to the standard protocol of the NEBNext Ultra RNA Library Prep Kit for Illumina (New England BioLabs). The synthesized double-stranded cDNA was end-repaired using NEBNext End Prep Enzyme Mix before ligation with NEBNext Adaptor for Illumina. After USER enzyme treatment, cDNA was amplified by PCR under the following conditions: 20 µL cDNA, 2.5 µL Index Primer, 2.5 µL Universal PCR Primer, and 25 µL NEBNext Q5 Hot Start HiFi PCR Master Mix 2X; 98 °C for 30□s; 12 cycles of 98□°C for 10□s and 65□°C for 75□s; and 65□°C for 5□min. The cDNA library was sequenced with a NextSeq 500 instrument (Illumina) using 150-bp paired-end reads with a NextSeq 500 High Output Kit (300 cycles).

### Genome assembly and contaminant elimination

#### Caerostris darwini genome

Long reads from Nanopore sequencing were first quality filtered using NanoFilt with the options -q 7 –headcrop 50 ^38^ and then assembled using Flye 2.7 ^39^ with three Racon polishing iterations ^40^. The assembly was further polished by Pilon v 1.23 ^41^ over three rounds using the Illumina sequencing reads of the 10X GemCode library. Synthetic long reads from 10X GemCode library sequencing were assembled by Supernova 2.1.1 in pseudohap2 output mode ^42^. Finally, the two assemblies were merged using quickmerge with the options -hco 5.0 -c 1.5 -l 100000 -ml 5000 ^43^.

#### Caerostris extrusa genome

Long reads from Nanopore sequencing were first quality filtered using NanoFilt with the options -q 7 –headcrop 50 ^38^ and then assembled using Flye 2.7 ^39^ with three Racon polishing iterations ^40^. The assembly was further polished by pilon v 1.23 ^41^ over three rounds using the Illumina sequencing reads of the cDNA library.

The contaminants in the genome assemblies were eliminated based on BlobTools analysis ^44^. The genome sequence was subjected to a Diamond ^45^ BLASTX search (--sensitive --max-target-seqs 1 --evalue 1e-25) against the UniProt Reference proteome database (downloaded on 2018 Nov.). The coverage data were calculated by sequence read mapping with BWA MEM (Burrows-Wheeler Alignment v0.7.12-r1039) ^46^. Contigs that were classified as bacterial, plant, or fungal sequences were removed from the assembly. Mitochondrial contigs were also removed. The filtered genome assembly was assayed by using BUSCO v4.0.5 (eukaryote lineage) ^20^ to validate genome completeness.

### Gene prediction and annotation

Genes were predicted using a gene model created from cDNA-seq mapping data. The cDNA-seq reads were mapped to the reference genome with HISAT2 (v2.1.0) ^47^. Repeat sequences were detected by RepeatModeler (1.0.11) and soft-masked by RepeatMasker (v4.0.7) (http://www.repeatmasker.org). The soft-masked genome was subjected to gene prediction with BRAKER (v2.1.4, --softmasking --gff3) ^47,48^. The numbers of predicted protein-coding genes were initially 56,145 and 82,821 for *C. darwini* and *C. extrusa*, respectively. The predicted genes were annotated by Diamond BLASTP searches against public databases (UniProt TrEMBL, UniProt Swiss-Prot). Redundant genes were eliminated by CD-HIT-EST ^49^ clustering with a nucleotide identity of 97%. Furthermore, we collected the genes with an expression level of more than 0.1 and annotated them to obtain functional gene sets. Finally, functional protein-coding gene sets of 16,907 and 18,655 genes were obtained. BUSCO (v4.0.5) was used to determine the quality of our functional gene set using the eukaryote lineage.

### Spidroin catalogue curation

Spidroin genes identified in the bark spiders were curated based on a Spidroin Motif Collection (SMoC) algorithm ^9,15,23,50,51^. This spidroin curation algorithm was implemented using the hybrid assembly of short and long reads. The de Bruijn graph assembly of Illumina short reads was used for N/C-terminal domain searching by homology searches. The obtained terminal domains were used as seeds for screening the short reads harbouring exact matches of extremely large k-mers extending to the 5′-end, and the short reads were aligned on the 3′-side of the matching k-mer to build a position weight matrix (PWM). Based on stringent thresholds, the terminal domains were extended until the next repeats appeared. Finally, the collected full-length subsets of the repeat units were mapped onto error-corrected Nanopore long reads. The data on spidroin gene length or architecture were curated manually based on the mapped long reads. Therefore, the full length could be obtained within one long-length read.

### Phylogenetic tree

The spidroin phylogenetic trees were constructed by MEGAX ^52^ based on the first 100 N-terminal amino acid residues of the corresponding available spidroin sequences in the family Araneidae. The collected spidroin genes were aligned with MUSCLE, and the phylogenetic relationships were calculated using NJ. FigTree version 1.4.3 (http://tree.bio.ed.ac.uk/software/figtree/) was used as the viewer for the trees.

### Proteome analysis

The proteome analysis of dragline silks was performed with nanoElute and timsTOF Pro (Bruker Daltonics, Bremen, Germany). The reeled dragline silks were immersed in 200 µL of lysis buffer [6 M guanidine-HCl, pH 8.5] per mg silk and frozen in liquid nitrogen. After 10 sonication cycles of 60 sec on/off at a high level using a Bioruptor II sonicator (BM Equipment), the protein content was quantified with a Pierce BCA Protein Assay Kit (Thermo Scientific). The lysate [50 µg protein/50 µL] was incubated with 0.5 µL of 1 M DTT solution at 37 °C for 30 min, followed by 2.5 µL of 1 M IAA solution at 37 °C for 30 min in the dark. After fivefold dilution with 50 mM NH4HCO3. The sample was acidified with TFA and desalted with C18-StageTips ^53^. A 200 ng aliquot of the digested sample was injected into a spray needle column [ACQUITY UPLC BEH C18, 1.7 µm, Waters, Milford, MA, 75 µm i.d.×250 mm] and separated by linear gradient elution with two mobile phases (A [0.1% formic acid in water] and B [0.1% formic acid in acetonitrile]) at a flow rate of 280 nL/min. The composition of mobile phase B was increased from 2% to 35% over 100 min, changed from 35% to 80% over 10 min and kept at 80% for 10 min. The separated peptides were ionized at 1600 V and analysed in a parallel accumulation serial fragmentation (PASEF) scan ^54^. Briefly, the PASEF scan was performed at ion mobility coefficients (1/K0) ranging from 0.6 Vs/cm2 to 1.6 Vs/cm2 over a ramp time of 100 msec, keeping the duty cycle at 100%. An MS scan was performed in the mass range of m/z 100 to m/z 1700, followed by 8 PASEF-MS/MS scans per cycle. Precursor ions were selected from Top N intense ions in a TIMS-MS survey scan (precursor ion charge: 0-5, intensity threshold: 1250, target intensity: 10000). In addition, a polygon filter was applied to the m/z and ion mobility plane to select the most likely representative peptide precursors without singly charged ions. CID was performed with the default settings (isolation width: 2 Th at m/z 700 and 3 Th at m/z 800, collision energy: 20 eV at 1/k0 0.6 Vs/cm2 and 59 eV at 1/k0 1.6 Vs/cm2). De novo sequencing and database searches were performed with an error tolerance of 20 ppm for precursor ions and 0.05 Da for fragment ions with PEAKS software. The protein sequence database generated from our draft genome was used for protein identification. The raw MS data and the analysis files have been uploaded to the ProteomeXchange Consortium from the jPOST partner repository ^55^ under accession number PXD026224. To eliminate the possibility that other silks might have contaminated our reeled dragline silk samples, we removed background data. We separated clearly the dragline and minor ampullate silks from the spinnerets by microscopy and subjected them to LC-MS analysis to identify the background proteins. Using the iBAQ score, we identified the proteins specifically expressed in the minor ampullate silk based on a significance level of FDR < 0.01 and removed them from the results of the dragline silk proteome analysis as contaminants.

### Gene expression analysis

mRNA expression profiling was conducted from the cDNA-seq data. Gene expression levels were quantified and normalized as transcripts per million (TPM) values by mapping the processed reads to our assembled draft genome references with Kallisto version 0.42.1^56^.

### Measurement of bark spider silk properties

The surface morphology of the dragline silks was observed by SEM (JCM 6000, JEOL Ltd., Tokyo Japan). Samples were mounted on an aluminium stub with conductive tape backing and sputter-coated with gold for 1 min using a Smart Coater (JEOL) prior to SEM visualization at 5 kV. At least 8 individual mechanical stretching tests were performed for each dragline silk. The experimental setup was similar to those reported previously ^57^. Each fibre was attached to a rectangular piece of cardboard with a 5 mm aperture using 95% cyanoacrylate. The tensile properties of the fibres were measured using an EZ-LX universal tester (Shimadzu, Kyoto, Japan) with a 1 N load cell at a strain rate of 10 mm/min (0.033 s-1) at 25 °C and 48% relative humidity. For each tensile test, the cross-sectional area of an adjacent section of the fibre was calculated based on the SEM images.

### Measurement of WAXS

The crystalline state of the dragline silks was measured by synchrotron WAXS analysis in the SPring-8 BL05XU beamline, Harima, Japan, according to a previous report ^58^. The X-ray energy was 12.4□keV at a wavelength of 0.1□nm. The sample-to-detector distance for the WAXS measurements was approximately 257□mm. The exposure time for each diffraction pattern was 10 s. The resultant data were converted into one-dimensional radial integration profiles using Fit2D software ^59^. The resultant data were corrected by subtracting the background scattering. The degree of crystallinity was evaluated from the area of the crystal peaks divided by the total area of the crystal peaks and the amorphous halo by fitting the Gaussian function using Igor Pro 6.3.

## Data availability

All raw reads and assembled sequence data have been uploaded to DDBJ under BioProject numbers PRJDB11546 and PRJDB11547. The whole-genome sequence is available at the Whole-Genome Shotgun (WGS) database of DDBJ under accession numbers BPLQ01000001-BPLQ01015733 (*C. darwini*) and BPLR01000001-BPLR01021729 (*C. extrusa*). The raw MS data and analysis files have been uploaded to the ProteomeXchange Consortium from the jPOST partner repository ^55^ under accession number PXD026224.

## Acknowledgements

The authors thank Akio Tanikawa for the morphological identification of spiders and the Malagasy Institute for the Conservation of Tropical Environments (MICET) (especially Tiana Vololontiana), the Ministry of Environment and Sustainable Development of Madagascar (Ministère de l’Environnement de l’Ecologie et des Forêts at that time), the Mention Zoologie et Biologie Animale (MZBA), Université d’Antananarivo, and Madagascar National Parks (MNP) for their cooperation and permission to conduct sampling in Madagascar. The authors also thank Yuki Takai and Naoko Ishii for technical support in the sequencing analysis and Sumiko Ohnuma for technical support in proteome analysis. The work was supported by a Nakatsuji Foresight Foundation Research Grant, the Sumitomo Foundation (190426), and a Grant-in-Aid for Scientific Research (B) (21H02210) to N.K., a grant from the ImPACT Program of the Council for Science, Technology and Innovation (Cabinet Office, Government of Japan) to H.N., K.N., and K.A., and a grant from the Yamagata Prefectural Government and Tsuruoka City, Japan, to N.K., M.M., and K.A.

## Author contributions

K.N. and K.A. designed the entire project. N.K. and K.A. performed transcriptome and genome sequencing and assembly, and conducted expression analysis. N.K. and M.M. performed proteome analysis. H.N., R.O., A.D.M., and K.N. collected spider and silk samples, examined mechanical properties, and provided photographs of *C. darwini* and *C. extrusa*. N.K. and Y.Y. analysed and curated the genome data. K.N. and H.M. performed the synchrotron experiments. N.K. and K.A. managed the computer resources. N.K. wrote the manuscript, and all authors contributed to editing and revising the manuscript.

