## Supplementary Fig. for "Darwin’s bark spider shares a spidroin repertoire with *Caerostris extrusa* but achieves extraordinary silk toughness through gene expression"

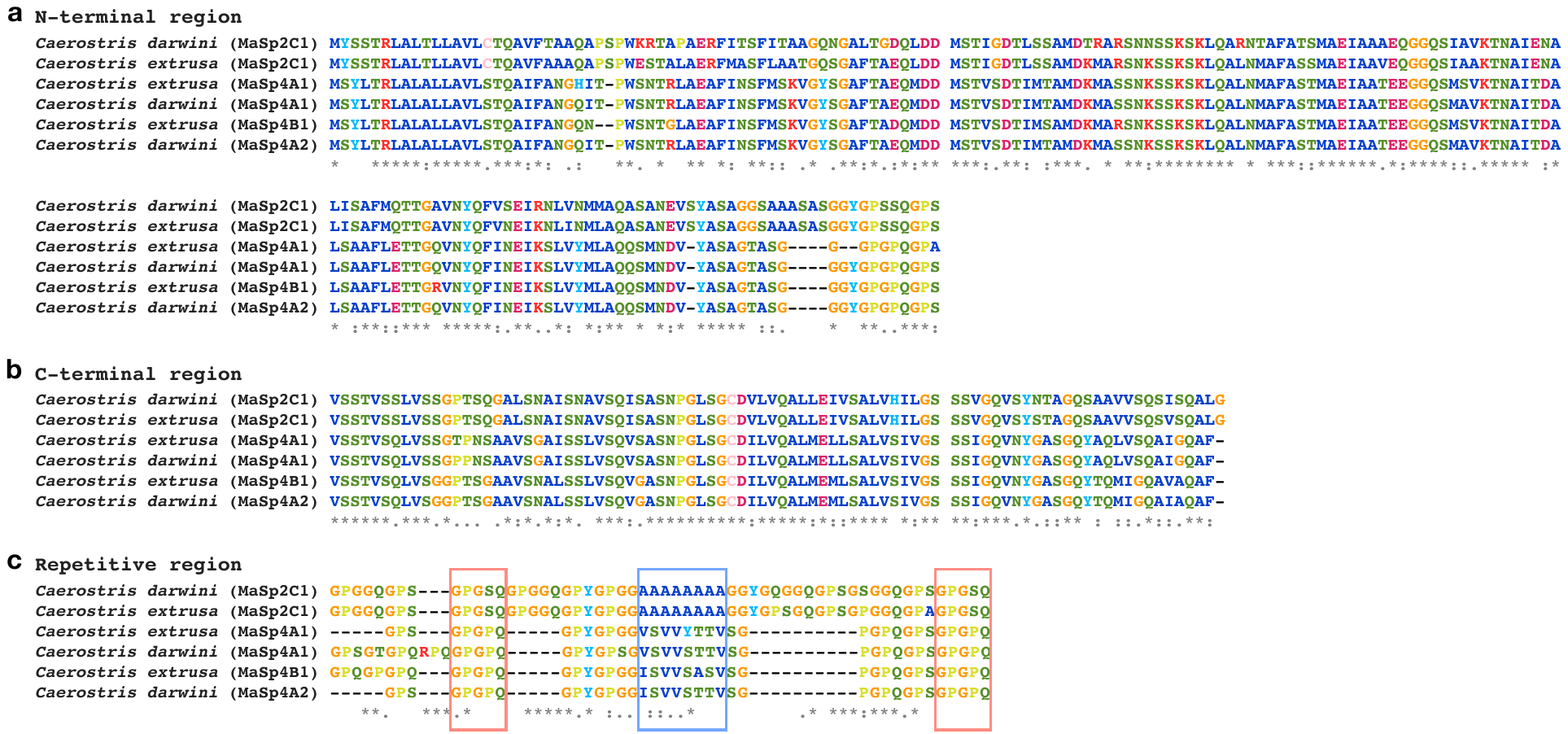

**Supplementary Figure 1. Alignment results of N/C-terminal and repetitive region of MaSp2 and MaSp4.**

This figures represent the alignment results of N-terminal (**a**), C-terminal (**b**), and repetitive (**c**) region of MaSp2 and MaSp4 in *C. darwini* and *C. extrusa*. The boxes at the repetitive region alignment mean the repeat body and tail (see Table 3).

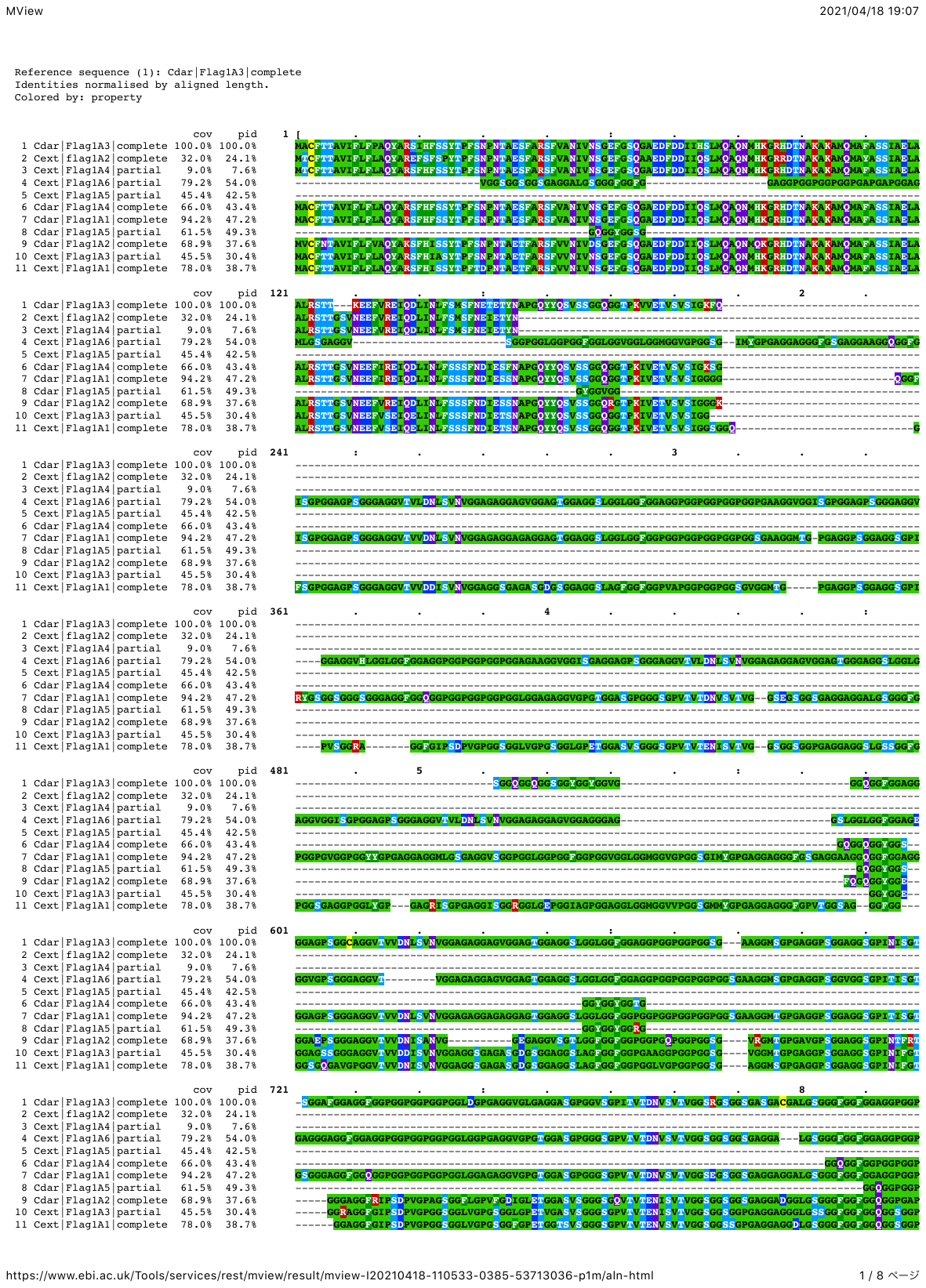

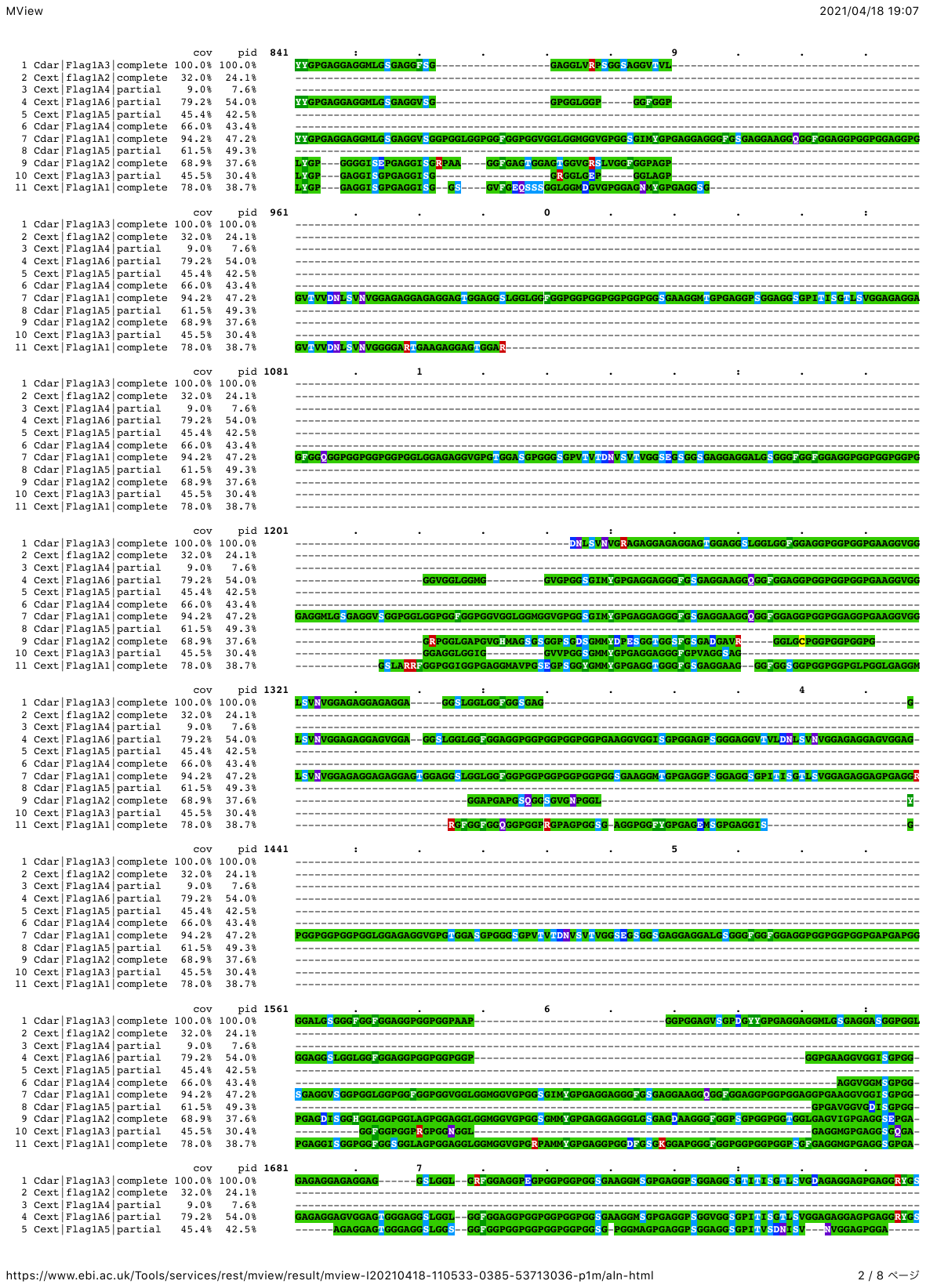

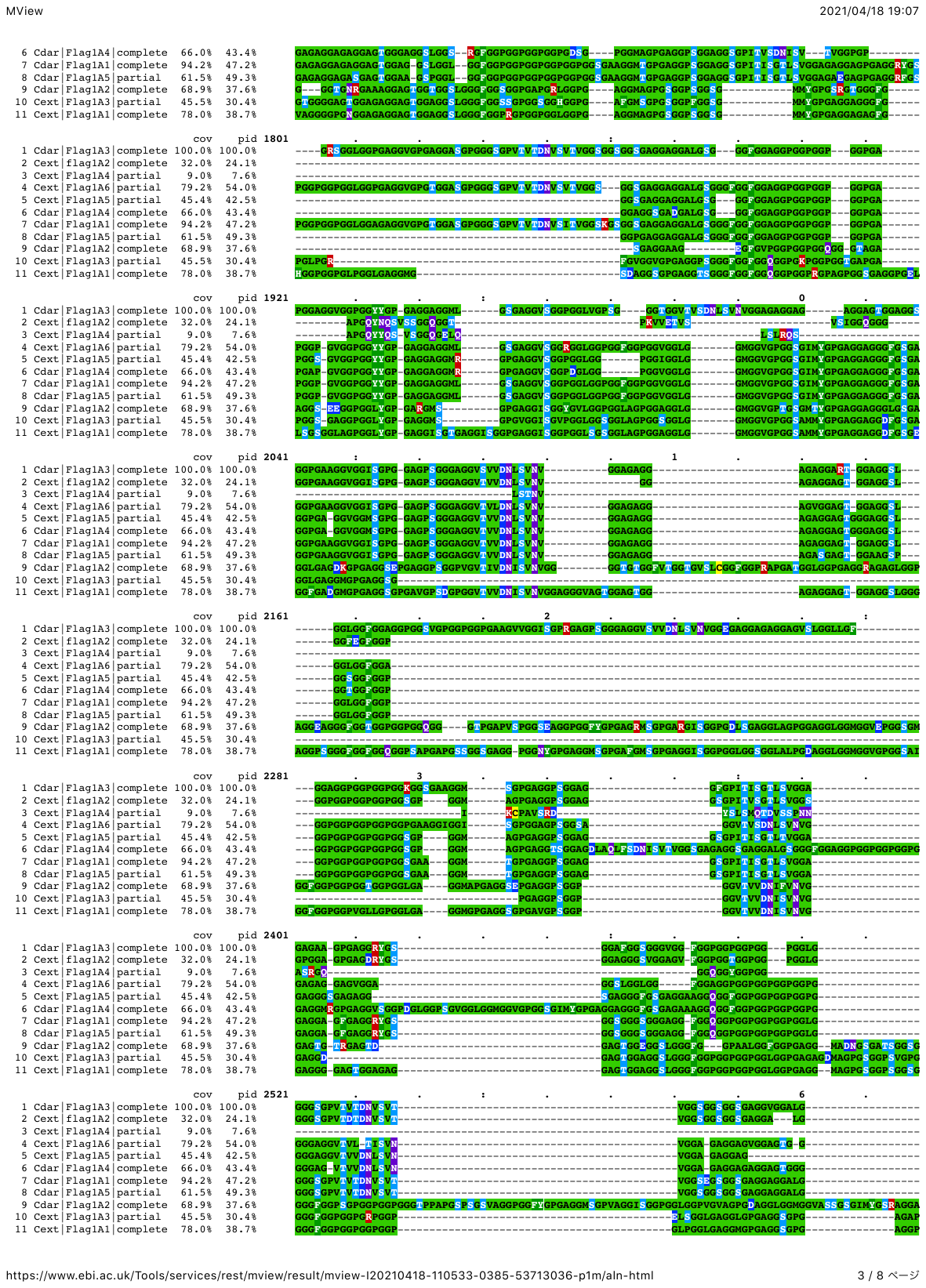

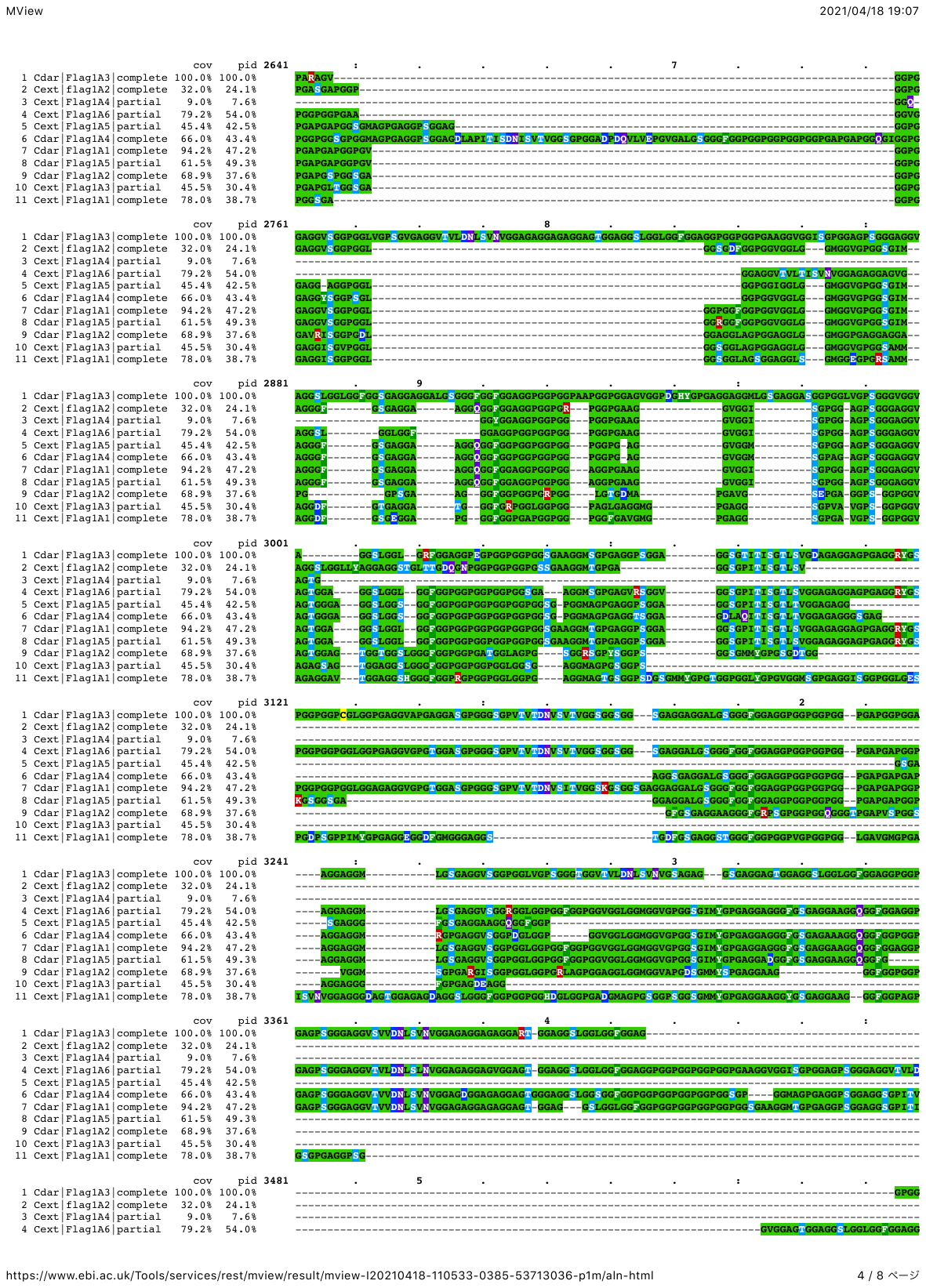

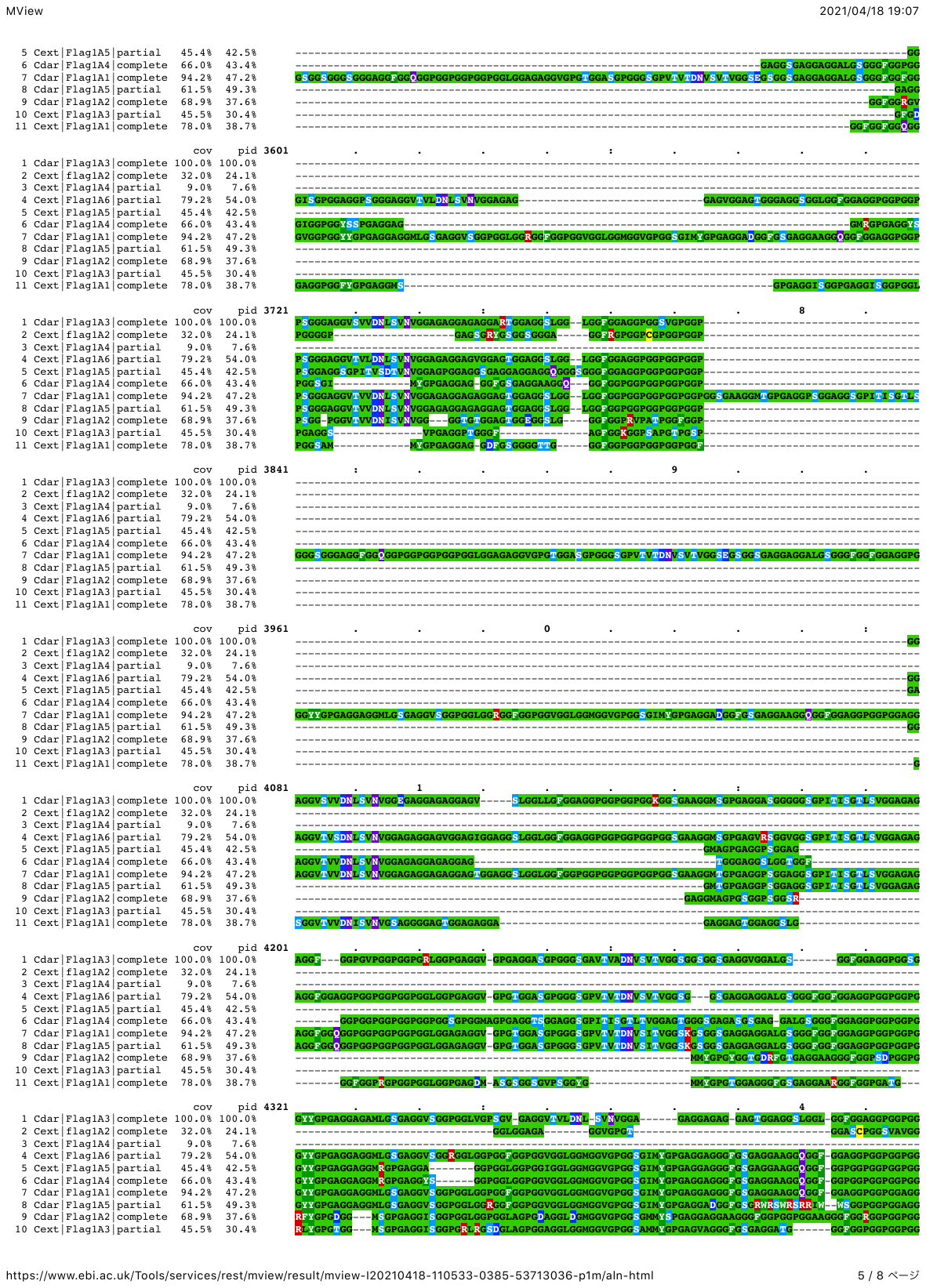

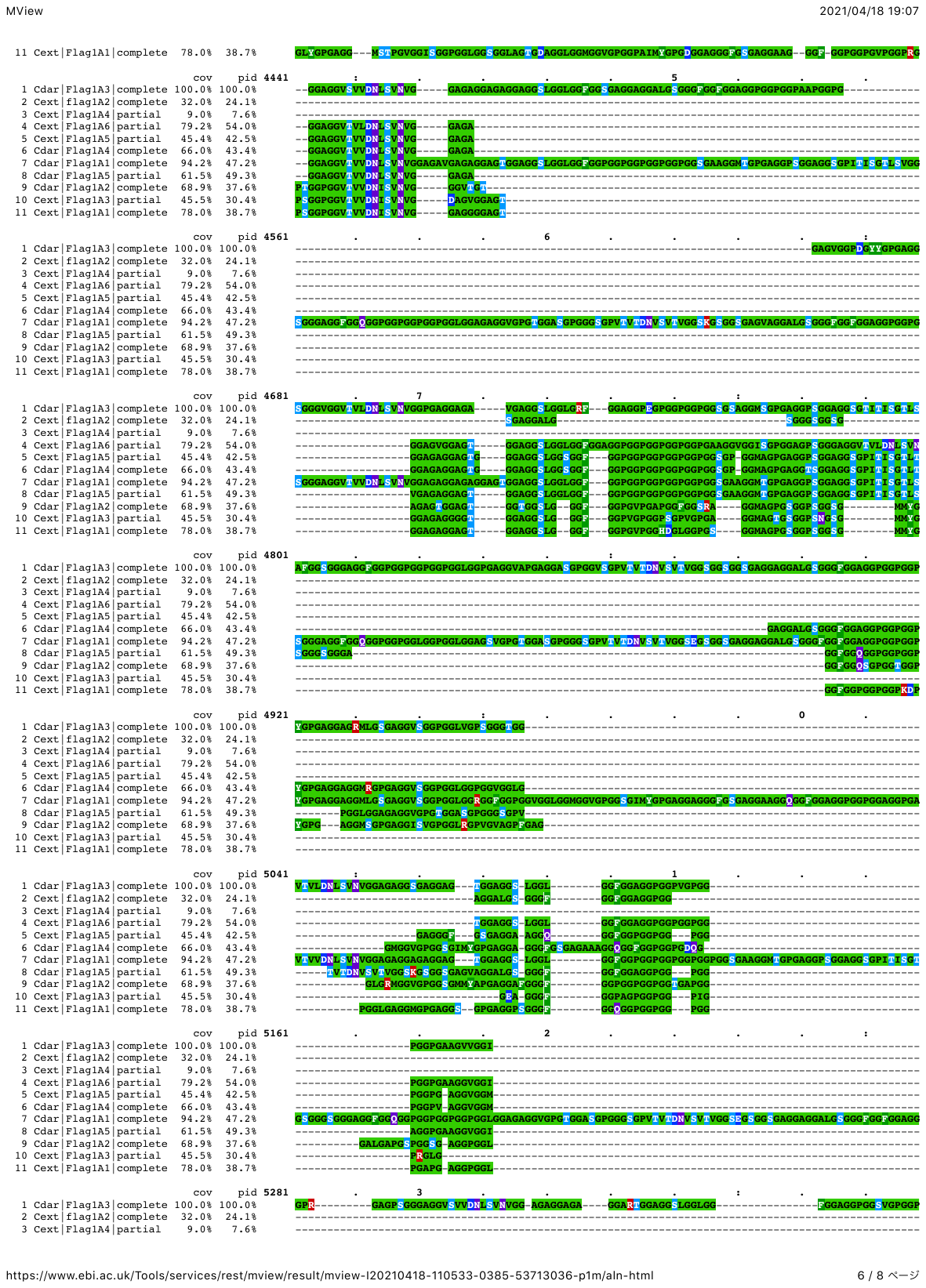

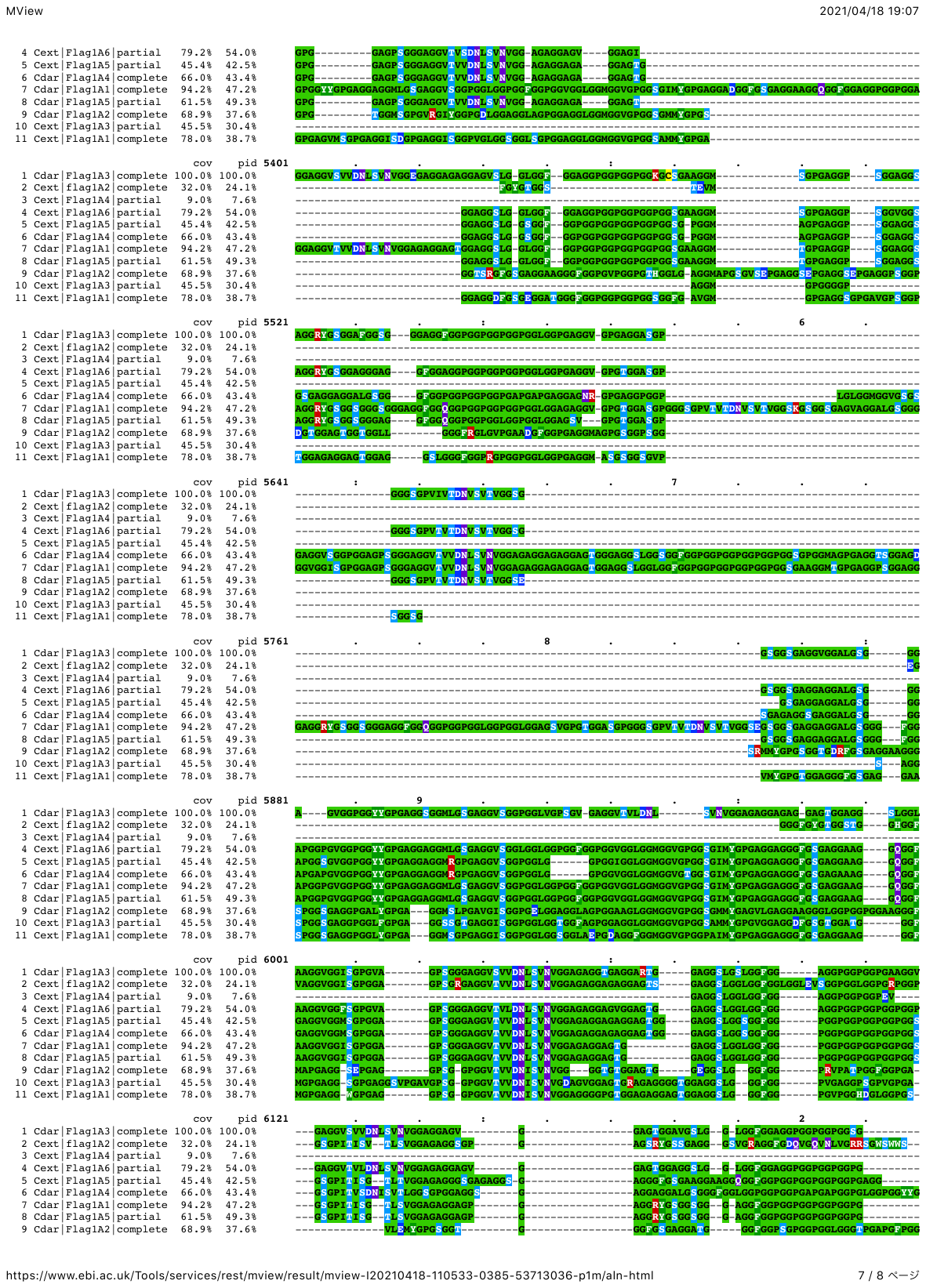

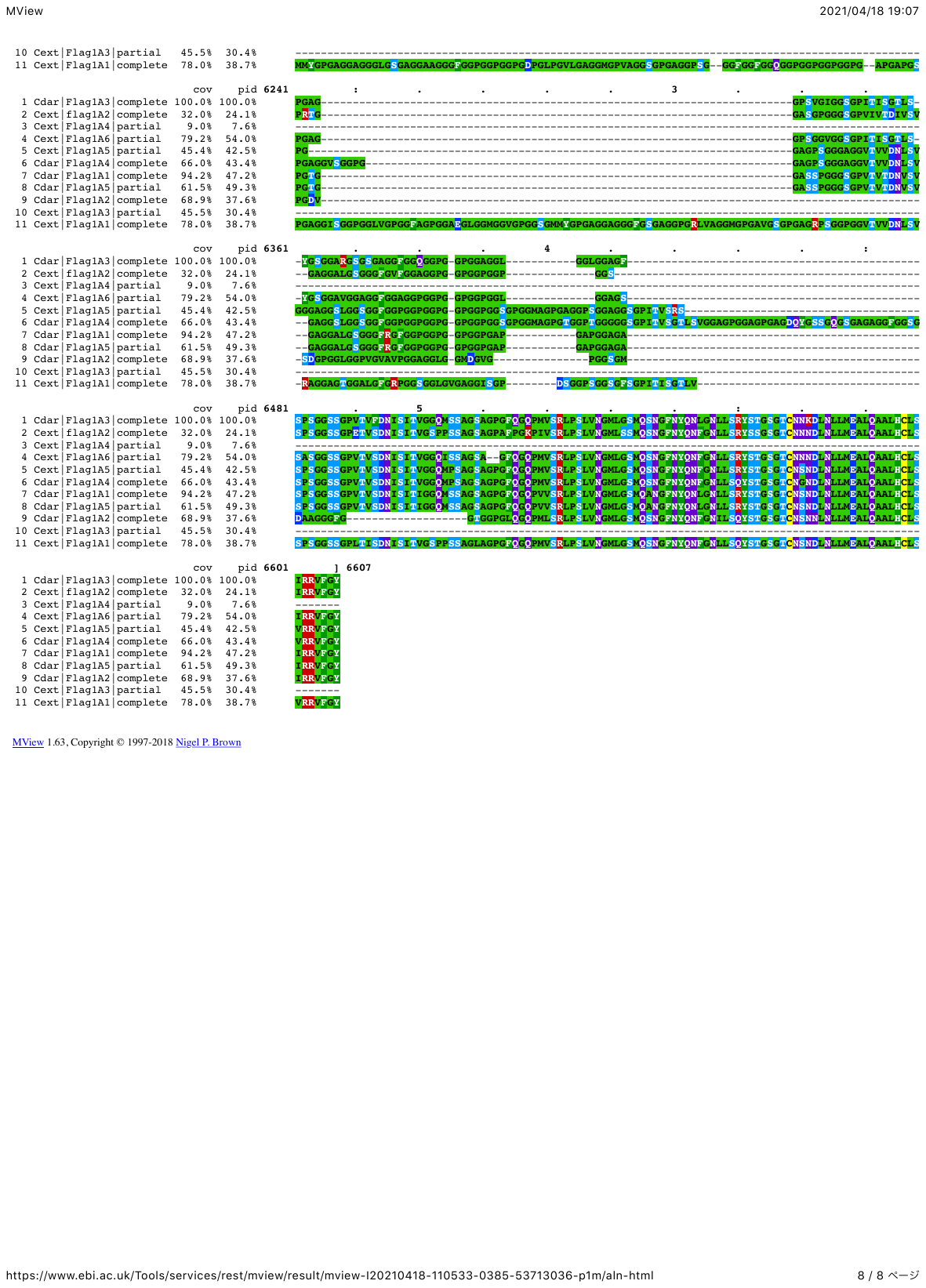

**Supplementary Figure 2. Alignment results of Flag amino acid sequence.**

Five and six Flag paralogues, including full-length (complete) and partial, were found in *C. darwini* (Cdar) and *C. extrusa* (Cext).

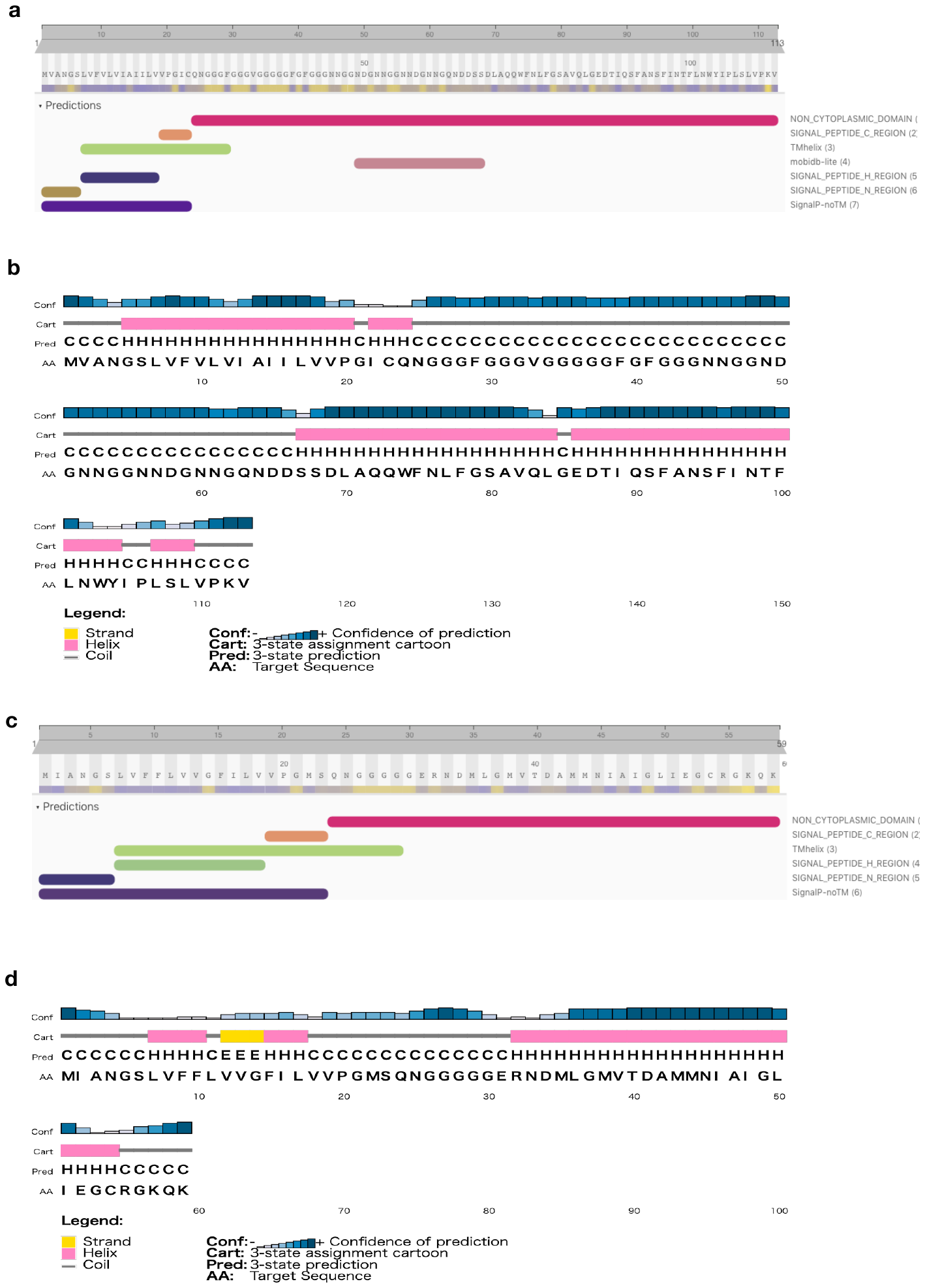

**Supplementary Figure 3. Secondary structure and motifs of SpiCE-CMa3.**

Secondary structure and motif of SpiCE-CMa3 (**a** and **b**: *C. darwini*, **c** and **e**: *C. extrusa*) were predicted by PSIPRED^1^ and InterProScan^2^.
